# Scalable spatial DNA sequencing from archival tissue maps copy number subclones

**DOI:** 10.64898/2026.08.24.746226

**Authors:** Cristian Soitu, Ugur Sahin, Anette Magnussen, Ashley Wong, Willem Bonnaffé, Mengran Fan, Merve Bilici, Simon Davis, Roman Fischer, Jasmine Reese, Toby House, Sorayya Moradi, Renuka Teague, Olaf Ansorge, Stefano Malacrino, Nasullah Alham, Emma McGregor, David Maldonado-Perez, Ian Tomlinson, David Wedge, Joanna Hester, Fadi Issa, Claire Edwards, Richard Bryant, Ian Mills, Jens Rittscher, Freddie Hamdy, Dan Woodcock, Clare Verrill, Srinivasa Rao

## Abstract

Spatially resolved DNA sequencing holds promise due to its potential utility in understanding cancer intra-tumour heterogeneity and tumour evolution in relation to tissue architecture. However, it has so far been used to a limited extent due to technical challenges and high cost of existing methods. Hence we aimed to develop a high throughput spatial genomic assay to obtain copy number alteration (CNA) information at user-defined spatial resolution.

We derived CNA profiles from ultra-low coverage whole genome sequencing at sub-millimetre resolution from archival samples using a novel method called Adaptive Resolution Multiscale Spatial DNA sequencing (ARMS DNAseq). We used it to profile CNAs from more than 766 regions (tiles) from 3 patients, covering a total area of over 300 mm^2^, with 1.2–2.6 million mapped reads per tile and tile sizes of 0.1-0.99mm^2^.

Using ARMS DNAseq, we delineate tumour evolution in a spatial context, and identify more tumour subclones that were obscured or incompletely represented in bulk multi-region whole genome sequencing. Next, we show associations between tumour subclones and morphology, and prediction of subclone identity from deep learning-derived image representations. Finally, we demonstrate multi-omic integration by alignment with spatial transcriptomic data, showing subclone-specific immune cell co-occurrence as well as transcriptional programmes cutting across subclone boundaries. ARMS DNAseq converts low-throughput, region-by-region profiling into a scalable and adaptable workflow for direct spatial copy number profiling from archival tissue sections.

## BACKGROUND

Sequencing DNA from cancer samples provides insights into the development of malignancies by identifying somatic mutations. Using these data, subclonal structure and cancer cell phylogeny can also be determined. Such genomic profiling has largely been limited to bulk DNA sequencing, resulting in the loss of spatial context. The addition of a spatial dimension can lead to a better understanding of the organisation of cancer subclones and their relationship with tissue composition. Multi-region bulk sequencing has attempted to address this to a limited extent by analysing multiple samples from the same tumour, in pioneering work by Gerlinger et al.^1^ and many others since then ^2–4^. This approach combined with deconvolution of subclones results in increased subclonal resolution. It has been pointed out that increasing sampling density from a tumour is more informative than sequencing a few samples at greater depth^5,6^. In studies that analysed a large number of samples from the same patient, a high degree of genomic heterogeneity was observed^7,8^. However, this is generally not feasible for scaling beyond a few patients due to the high effort and costs involved in sampling, nucleic acids extraction, library preparation, and sequencing.

Previous work by us and others using bulk sequencing incorporated spatial information for the sampled regions, which revealed routes of tumour spread^9,10^. Recently developed spatial DNA methods such as Slide-DNAseq^11^, BaSISS^12^, and laser capture microdissection^13^ have been shown to work well for genomic profiling from small tissue sections, but the cost and effort is prohibitive when scaled up to a larger tissue area. In addition, many of these methods require fresh frozen tissue, which poses a significant barrier to sample access and experimental logistics. While spatial transcriptomics data have been used to infer copy number alterations^14^, it is an indirect method that is limited by genomic resolution (in the megabase range), variable accuracy, and reference requirements^15,16^. Hence, a gap exists for a high-throughput method to profile genomic DNA directly at tuneable genomic and spatial resolution from formalin fixed paraffin embedded (FFPE) tissues.

We have developed a new ‘mini-bulk’ method called Adaptive Resolution Multiscale Spatial DNA sequencing (ARMS DNAseq) to address this need. Our method scales up shallow whole genome sequencing to hundreds of regions of adaptable size, enabling us to profile large areas of FFPE sections from prostate cancer patients.

We demonstrate that ARMS DNAseq can be used to profile hundreds of regions from cancer tissue sections and identify spatially-distinct subclones at variable spatial resolution. We then use this method to test an association between genomic and morphological heterogeneity and show how it can be exploited to predict genomic subclones in whole slide images. We also show that ARMS DNAseq enables the annotation of tumour subclone phylogenies with morphological subtypes, thus allowing us to distinguish evolutionarily ‘early’ and ‘late’ morphologies. Finally, we show that ARMS DNAseq can be integrated with other spatial omics methods such as 10X Xenium, to understand subclone-specific gene expression programmes in relation to their respective microenvironments.

## RESULTS

### Spatial DNA sequencing workflow

Starting from FFPE sections (4-10 µm thickness), we scaled up sample throughput by semi-automation of the laser capture step. Firstly, we developed a Napari-based^17^ software tool called ARMS-tiler that takes region of interest annotations (provided by a pathologist or derived from upstream image analysis) as input and generates ‘tiles’ of user-selected shapes and sizes (Fig. 1A). Secondly, we developed a workflow to laser capture hundreds of tiles into multi-well plates, while maintaining a link between spatial coordinates and well ID (Fig. 1B). Thirdly, we employed a multi-well approach to parallelise the initial library preparation steps. We further improved throughput by adopting a ‘one-pot’ and early pooling approach, which involves well-based adapter barcoding followed by post-ligation pooling and sequencing (Fig. 1C). Finally, by significantly modifying the low input laser capture WGS method originally described by Ellis et al.^13^, we miniaturised reaction volumes. Short read sequencing data thus obtained can be processed using the ARMS R package to generate copy number heatmaps and spatial subclone maps (Fig. 1D). Notably, our protocol does not involve whole genome pre-amplification^18^ despite low input.

**Figure 1:**
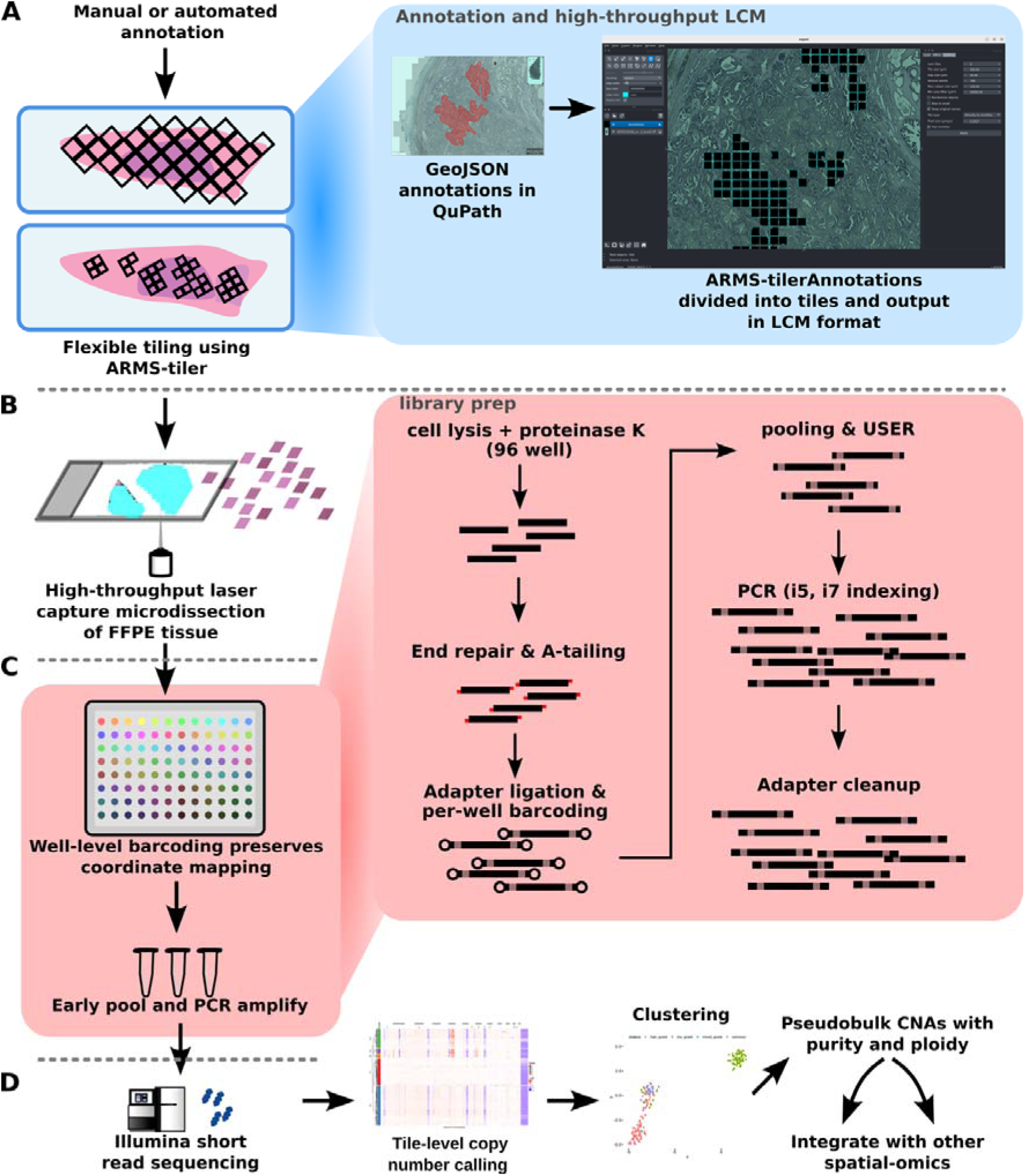
ARMS DNAseq workflow. A) Annotations (from upstream analysis or from pathologist) are divided into ‘tiles’ of user-defined spatial resolution using the ARMS-tiler software. B) Tiles are laser captured in a semi-automated process into 96-well plates, with a manifest linking well IDs to spatial coordinates. C) Whole genome library preparation is performed in the 96-well format until the ligation step, at which point adapters with well-specific barcodes are used. This enables post-ligation pooling, resulting in a single pooled sample per 96-well plate. Each pooled sample is further uniquely indexed at the PCR step to enable multiplexing. D) After Illumina short read sequencing, samples are demultiplexed at the plate and well level, to obtain reads mapping to each tile. These reads are then aligned to the genome and further processed downstream for copy number and subclone analysis.

### Technical performance of ARMS DNAseq

Two versions of ARMS DNAseq were used on three patient samples from the ProMOTE study^19^, of which P15 and P10 (also referred to as patients #15 and #10) were previously studied by us with bulk WGS^10^. Version A (which uses 10ul out of 40ul lysate and included a bead cleanup after lysis) was used for patient samples P15 and P10, whereas version B (which uses 5ul out of 40ul lysate as input and substituted bead cleanup for sample drying on the thermal cycler) was used for 10Q. While version B resulted in a 15% reduction in hands-on time, a higher percentage of duplicates were seen in this variation.

**Extended Table ET1:**
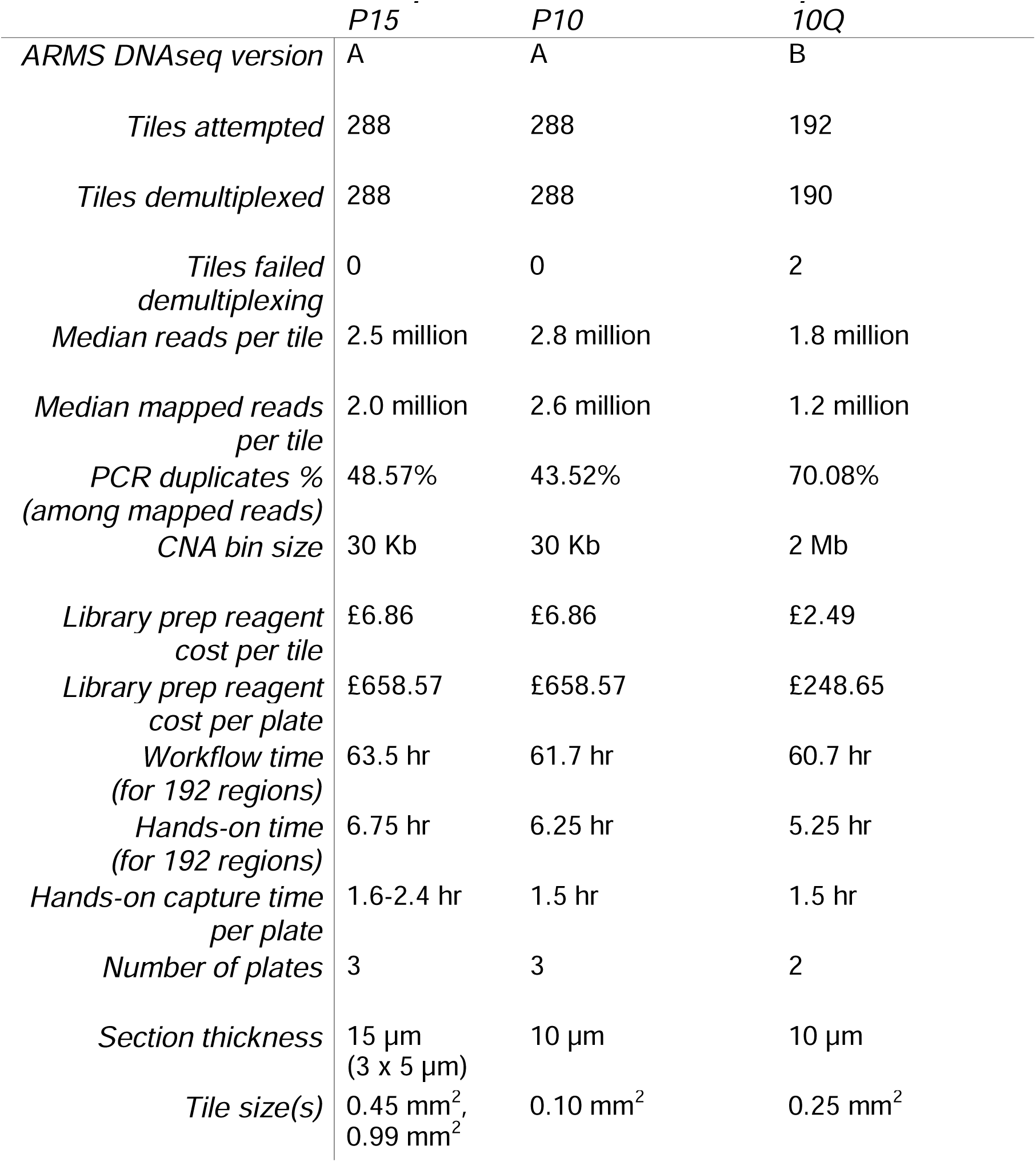
Technical performance of ARMS DNAseq.

### ARMS tiling maps CNA subclones across large FFPE tissue areas

As a first application of the ARMS DNAseq method, we sought to profile the majority of the tumour in a tissue section, while avoiding immune cell-rich areas in order to reduce reads from non-cancer cells. We selected three tissue sections from a tumour that we previously characterised extensively^10^, sampling close to previously profiled regions (Fig. 2A). Nearly the entire tissue area was selected, only excluding regions with high immune cell infiltration (determined by H&E), resulting in a total of 288 square tiles of either 0.99 or 0.45 mm^2^ area. The total tissue area profiled in this patient was approximately 233 mm^2^. Copy number alterations from these tiles are depicted as a heatmap with hierarchical agglomerative clustering used to group similar profiles (Fig 2B). This analysis revealed groups with distinct copy number profiles, which are termed hereon as copy number subclones or briefly as subclones. Visualisation of the copy number profile clusters by uniform manifold approximation and projection (UMAP) showed well-separated groups representing the two major cancer subclones (clusters 2 and 3) (Fig. 2C). Two additional subclones with distinct copy number profiles in the heatmap (clusters 5 and 8) are less distinct in the UMAP visualisation. In order to improve genomic resolution, data from all tiles within a subclone were combined to produce pseudobulk CNAs that revealed extensive multi-state oscillations in copy number in subclone 3 (Fig. 2D) consistent with the high number of inter-chromosomal translocations observed in bulk WGS in our previous work^10^. As the tiles were spatially barcoded during the library preparation, the corresponding subclone annotation could be mapped back to the tissue coordinates, revealing discrete spatial organisation of the subclones (Fig. 2E).

**Figure 2:**
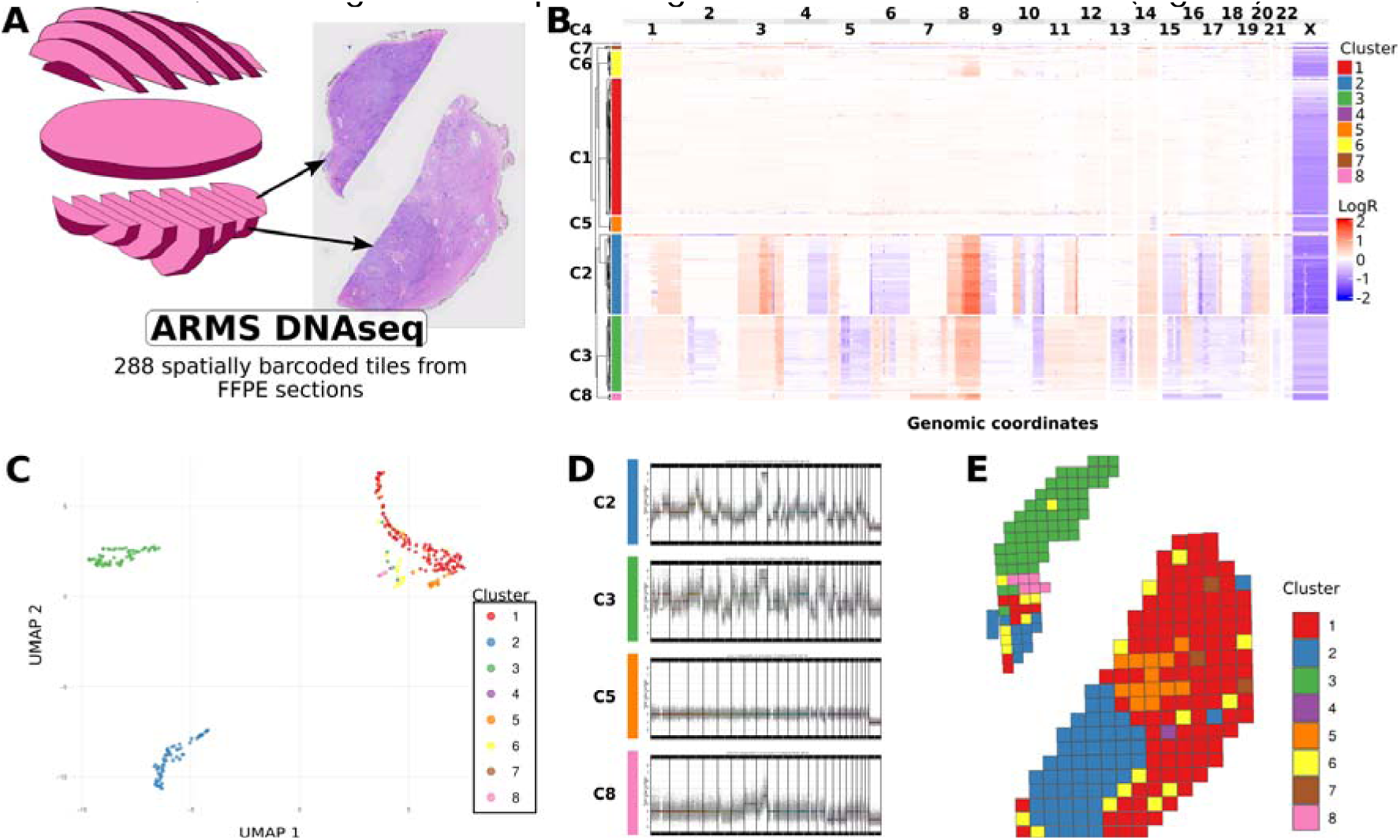
ARMS DNAseq in a radical prostatectomy FFPE sample (P15). A) A schematic shows the pathology blocks biobanked from this radical prostatectomy sample, with the arrows showing the specific block from which the tissue sections were obtained. B) Heatmap depicting Copy Number Alterations from each tile (rows) at each 30kb genomic locus (columns). C) Uniform Manifold Approximation and Projection (UMAP) visualisation of CNA data, showing distinct clusters. D) Pseudobulk analysis of each subclone (colours representing subclone ID). E) Spatial mapping of genomic subclones (colours representing subclone ID), with depicted tile size proportional to captured regions (0.45 mm^2^ tiles in the top-left tissue region, and 0.99 mm^2^ tiles in the bottom-right region).

Although spatial genomic profiling of the entire tumour area gives a more comprehensive understanding of subclone distribution, in some cases this may provide only marginal gains in information while resulting in higher costs. Hence, we adopted a modified strategy in another previously well-characterised patient sample. In this tissue section, we manually selected non-contiguous tumour areas with distinct histopathological appearances (in addition to some non-cancer regions) from a single tissue section and applied a random tiling strategy to sample only a proportion of the selected regions and avoiding most regions deemed to be non-informative (benign tissue, and regions with high immune cell content). As this resulted in a smaller total area, we could cover a larger span of this tissue section at a greater spatial resolution (tile size: 0.1 mm^2^). ARMS DNAseq identified tile-level copy number alterations, clustered into subclones (Extended Fig. E1 A). The tiles with their respective subclone identifier were subsequently mapped back to their tissue coordinates, showing some spatial demarcation of subclones, extensive heterogeneity and intermixing of subclones, and also non-contiguous distribution of subclones (e.g. C2, C3, C4) (Extended Fig. E1 B). The copy number profiles identified by ARMS DNAseq were also confirmed by our previously reported bulk WGS data from different regions of the same prostate (Extended Fig. E1 C). Furthermore, we discovered additional CNAs (e.g. 1q and 2p gains) through ARMS DNAseq which were not detectable in bulk WGS data (Extended Fig. E1 A, C).

**Extended Figure E1:**
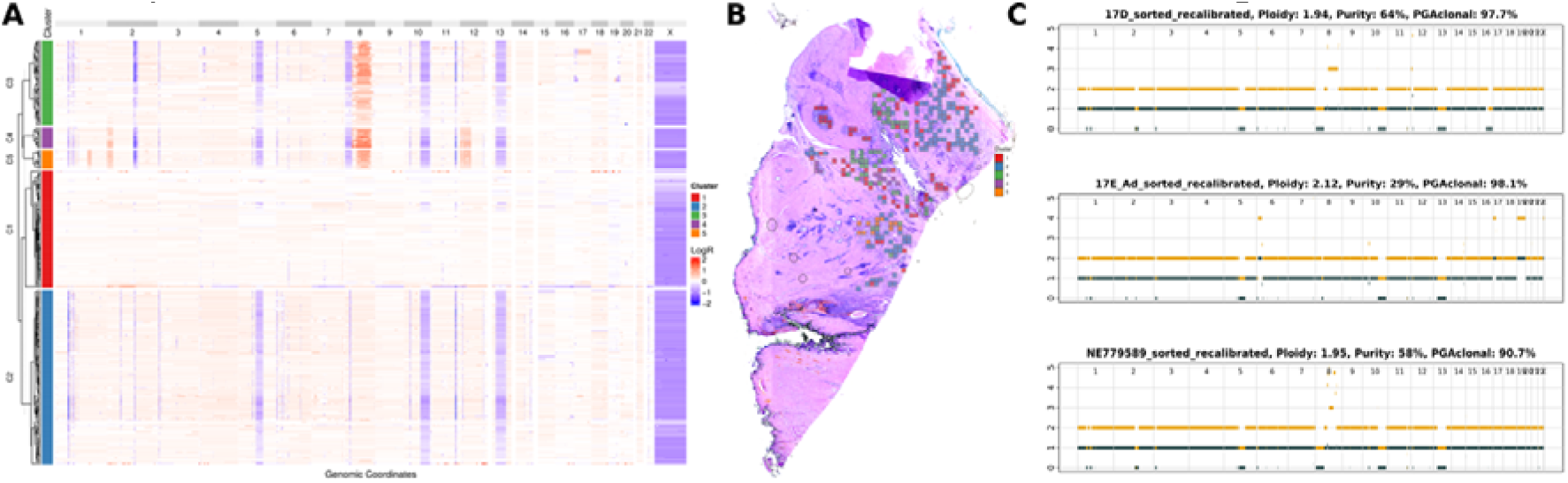
ARMS DNAseq in a second radical prostatectomy FFPE sample (P10). A) Heatmap depicting CNAs from tiles of approximately 0.1 mm^2^ area. B) Spatial mapping of genomic subclones (tile colours representing subclone ID) onto the whole slide image. C) CNA plots derived from bulk sequencing of fresh frozen tissue from adjacent blocks (top, middle) and lymph node samples (bottom), corroborating the CNAs seen with ARMS DNAseq.

**Extended Figure E2:**
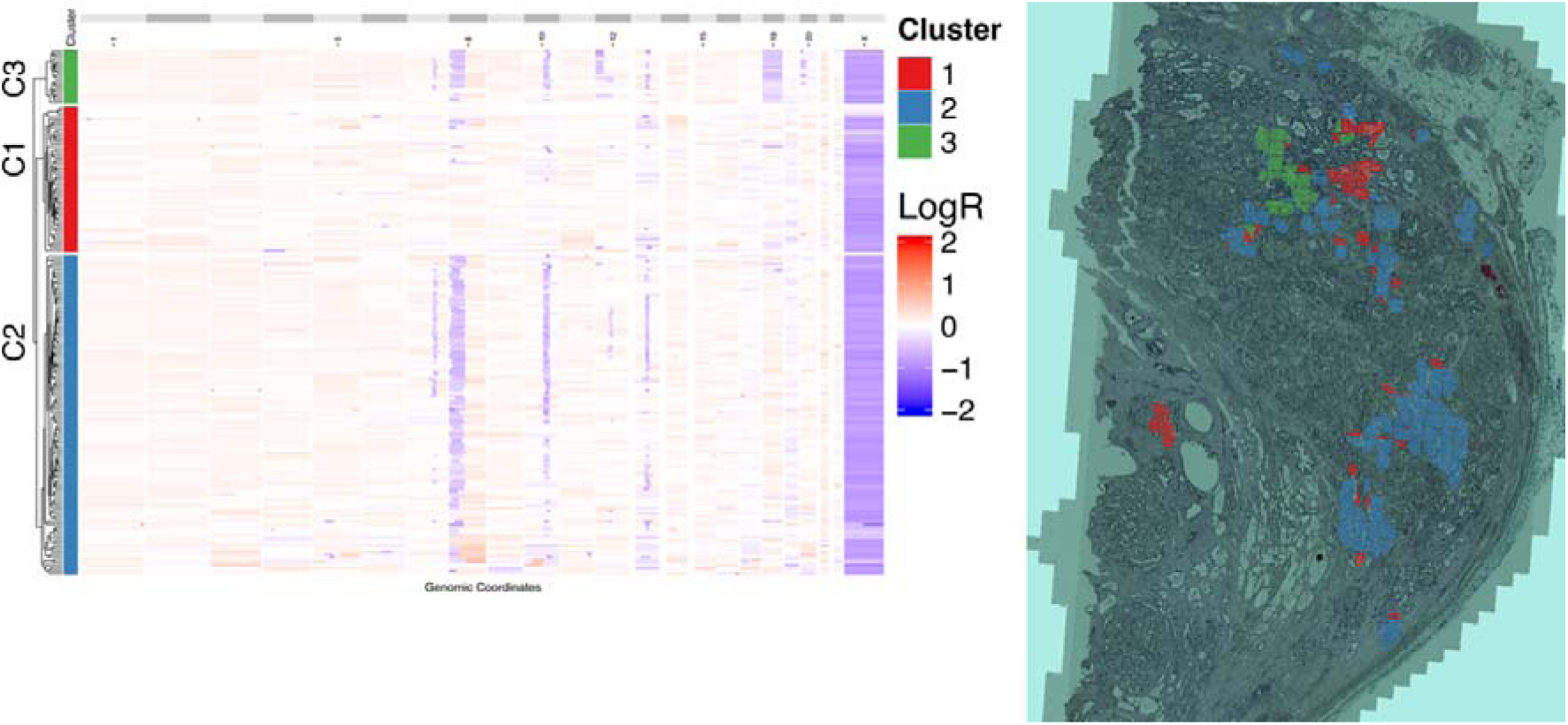
ARMS DNAseq in a third radical prostatectomy FFPE sample (10Q). A) Heatmap depicting CNAs from tiles of approximately 0.25 mm^2^ area. B) Spatial mapping of genomic subclones (tile colours representing subclone ID) onto the whole slide image.

### ARMS-derived subclone maps provide a genomic framework for morphological heterogeneity

Access to spatially resolved genomic data provided us with an opportunity to study the genotype-phenotype relationship. Histopathological analysis of the tissue areas corresponding to distinct subclones revealed that they have a morphology distinct from the other subclones (Fig. 3A). As visual distinction of histopathology is subjective, we sought to find an objective method by which morphology can be quantified. We extracted numerical representations (embeddings) of unannotated image patches using a foundational deep learning pathology model (Phikon v2) and performed hierarchical clustering on them to classify tissue morphologies (which we called morphology clusters; Fig. 3B). This unsupervised classification revealed differences in tissue architecture corresponding to biologically meaningful annotations such as stroma, immune rich regions, and Gleason patterns GP3, GP4 and GP5. More interestingly, a remarkable similarity was observed between morphological subtypes of malignant epithelium (morphology cluster 0: subclone C2; morphology cluster 5: subclone C3; morphology cluster 10: subclone C5) (Fig. 3C). A UMAP projection of the morphology embeddings (coloured by morphology cluster ID) with subclone identifiers overlaid on them shows the extent of this association (Fig. 3D). A map comparison analysis between the spatial distribution of genomic subclones and morphology clusters revealed statistically significant subclone-morphology associations (Fig. 3E).

**Figure 3:**
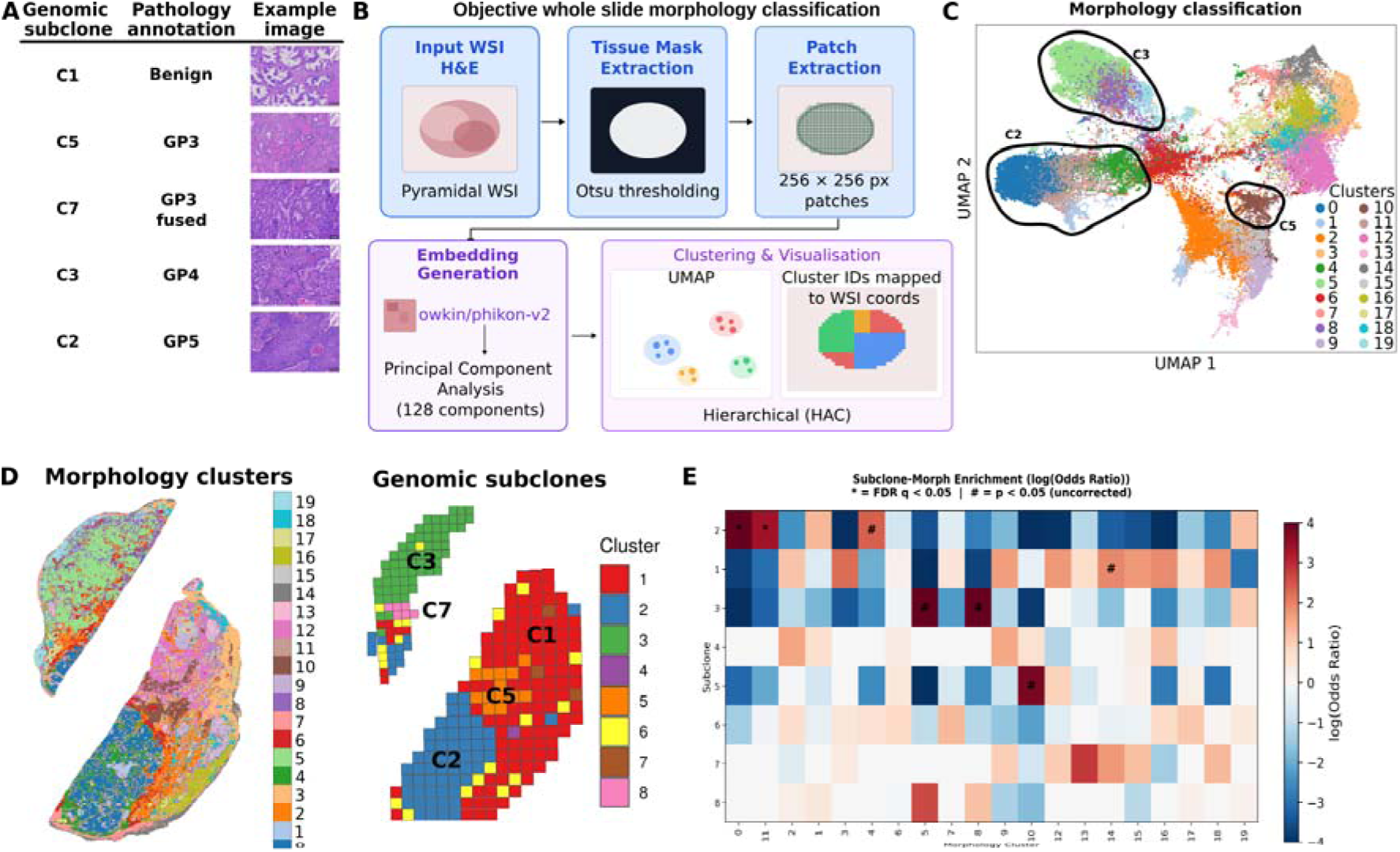
Spatial distribution of genomic subclones is significantly associated with morphology subtypes. A) Representative images of histology corresponding to each genomic subclone, with their respective pathological annotations. B) Workflow for extracting quantitative morphological representations of histopathological images using Phikon 2.0, a foundational pathology model. C) Comparison of the spatial distribution of objective morphological subtypes derived from Phikon 2.0 (left) and genomic subclones derived from ARMS DNAseq (right). D) UMAP of morphological embeddings from Phikon 2.0 (each dot representing a 256x256 pixel or 67.25x67.25 µm patch from the whole slide image) with colours representing morphological clusters, and black polygons overlaid to indicate the respective spatially overlapping genomic subclones. E) Association heatmap showing positive (red) or negative (blue) association between morphology cluster (X axis) and genomic subclone (Y axis).

### Morphology embeddings recover ARMS-defined genomic subclone identity across non-serial samples from the same patient

Having established that there is a strong association between genomic subclone identity and tissue morphology, we hypothesised that the former can be predicted from the latter. We trained a random forest model with genomic subclone labels and morphology embeddings (Fig. 4A), using 40% of the data (morphology patches) for training (Fig. 4B). The highest prediction accuracy was seen in the most abundant clones (clones C2 and C3; Fig. 4C). The model was applied to the entire sample (including the training subset), producing spatially coherent predicted regions going beyond the spatial resolution of ARMS DNAseq (Fig. 4D). Furthermore, we used the model to generate subclone predictions in a nearby tumour tissue from the same patient (transverse section; Fig. 4E). These predictions are validated by bulk WGS results (Fig. 4F), with the copy number profiles closely matching pseudobulk CNA calls from ARMS DNAseq (Fig. 2D). Extending the subclone prediction to all prostate sections from this patient enabled a 3D visualisation of cancer subclones across the entire prostate gland (Supplementary information 3).

**Figure 4:**
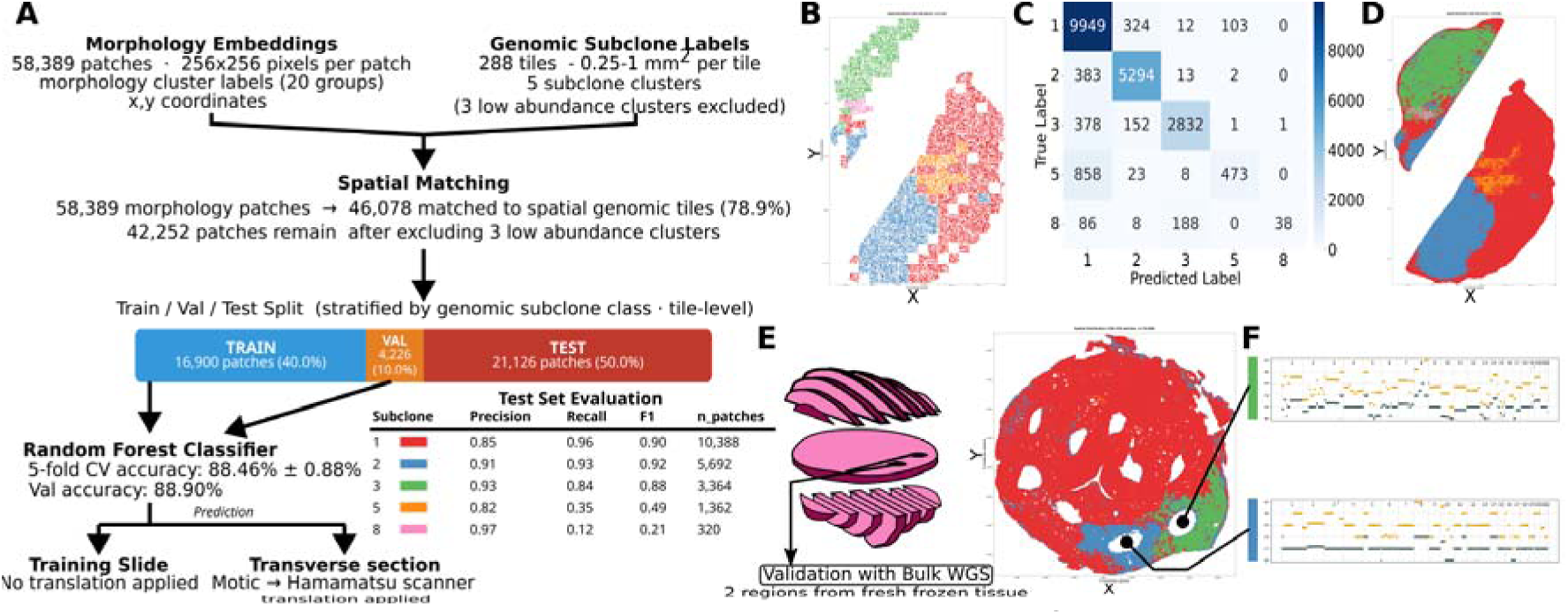
Tumour subclones can be predicted from morphology across samples within the same patient (patient #15). A) A workflow schematic showing how pathology foundation model-derived morphology embeddings and ARMS DNAseq subclone labels were used to train a random forest model. Subclone annotation colours shown in the Test Set Evaluation table apply to panels B, D, E, and F. B) 40% of the morphology patches from Fig. 2A were used for training, 10% for validation, and 50% were held out. C) Heatmap showing predicted vs. true labels, with the best performance for clusters IDs 1, 2, and 3. D) The random forest model was applied to the entire sample (including the original training set), revealing tissue contours at the resolution of the morphology patches. E) The random forest model was applied to an independent tissue slice, predicting two main genomic subclones (blue and green regions). F) Copy number analysis of the corresponding regions (green region = top; blue region = bottom) using traditional bulk sequencing closely matches the corresponding pseudobulk profiles from ARMS DNAseq shown in Fig. 2D.

### ARMS DNAseq enables identification of ‘early’ and ‘late’ morphologies

Phylogenetic reconstruction of the subclones identified by ARMS DNAseq in three patient samples reveals extensive copy number alteration driven tumour evolution (Fig. 5A, 5B, 5C). Morphologies (histopathological subtypes) can be placed on a phylogenetic tree, leading to the identification of evolutionarily early and late morphologies in these patients. We observed that in these samples, the early morphologies corresponded to low risk histopathological subtypes (Gleason Pattern 3) whereas the higher risk Gleason Pattern 4 and 5 subtypes occurred later in the cancer’s evolution.

**Figure 5:**
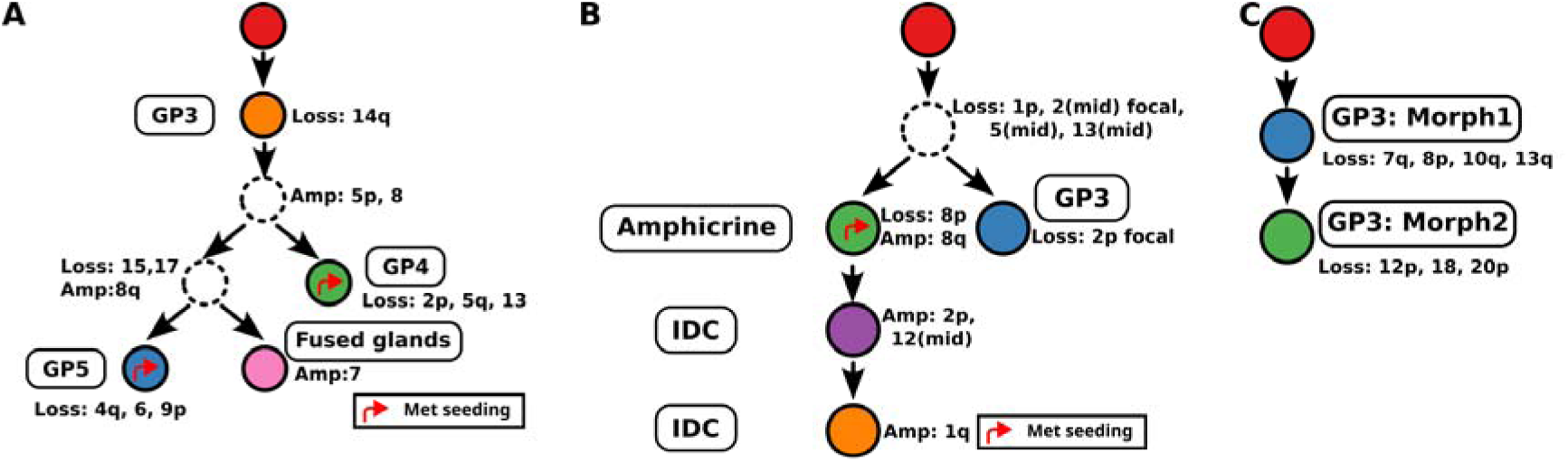
Phylogenetic trees depicting subclone evolution and associated morphologies. Phylogenetic analysis of three prostate cancer patients using ARMS DNAseq. The metastatic clone(s) (identified by matched bulk WGS-derived CNAs) are marked with a red arrow. Nodes in dotted lines indicate inferred subclones that were not found in the analysed tissue section. The node colours match the respective spatial tile colours shown in Figures 2E, E1B, and E2B.

### ARMS provides CNA ‘ground truth’ for spatial transcriptomic integration

While spatial genomics provides new insights into intra-tumour heterogeneity, tumour evolution, and genotype-phenotype relationships, DNA is only a part of the cancer puzzle. Integration of subclone information with other spatial-omics data can enable a more comprehensive understanding of the biological processes involved. Hence, we integrated single cell-level spatial transcriptomics data with ARMS DNAseq-derived subclone information (Fig. 6A). Based on the subclonal heterogeneity observed in the ARMS DNAseq data, a portion of the tissue was assayed with the Xenium platform (5100 gene panel). Regions of interest (Region 1 and Region 2) were outlined based on subclone predictions from H&E images (Extended figure E3).

**Extended figure E3:**
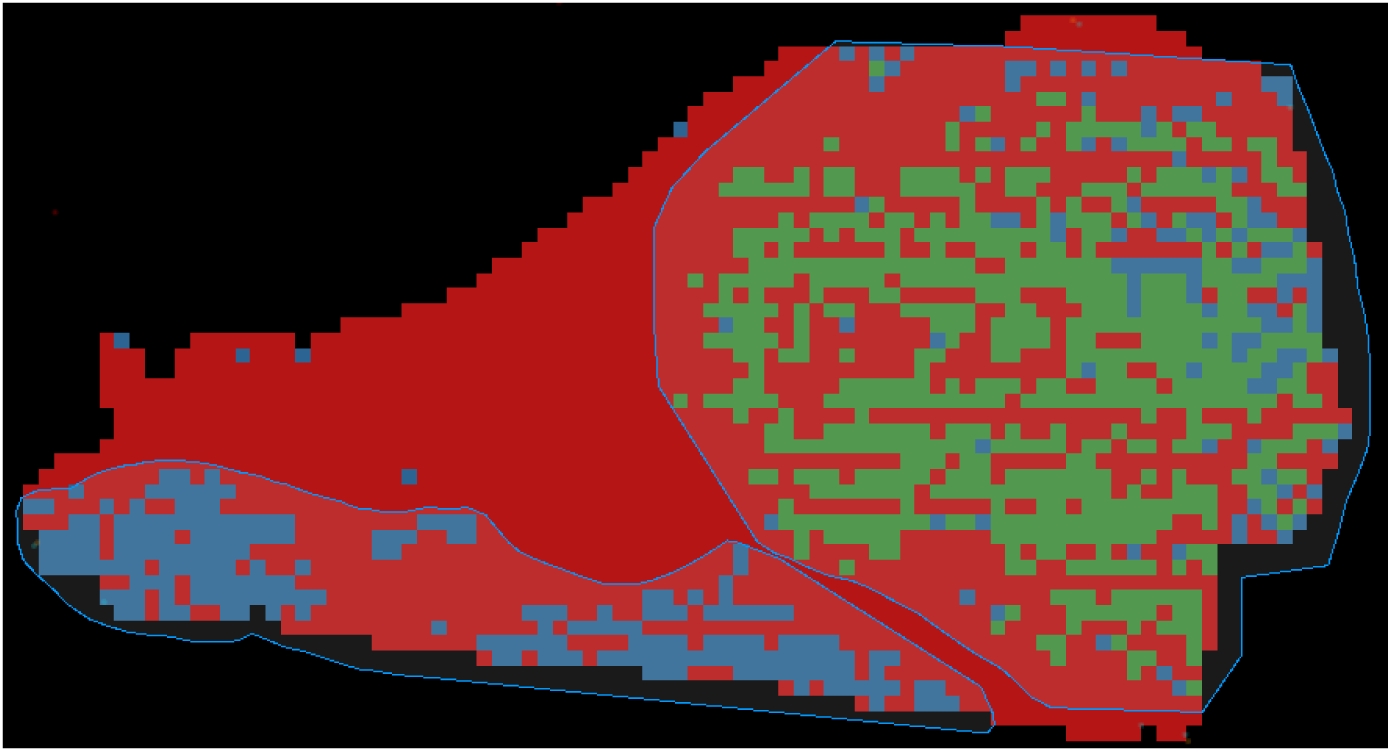
Morphology-based subclone prediction in a new tissue section used for Xenium spatial transcriptomic analysis, showing the predicted distribution of C2 (blue) and C3 (green). These predictions were used as the basis for outlining Region 1 (blue line, bottom) and Region 2 (blue line, right) in Fig 6B.

This integrative analysis showed co-localisation of distinct transcriptomic clusters with specific genomic subclones (Fig. 6B). Region 1 (corresponding to subclone C2) was seen to almost exclusively consist of cells belonging to the PCa 3 transcriptomic cluster, whereas Region 2 (subclone C3) consisted of three transcriptomic clusters PCa 1, PCa 2, and PCa 4 (Fig. 6C). The subclone distribution was largely mutually exclusive. Region 1 and Region 2 showed distinct gene expression patterns, with differences prostate epithelial markers (KLK2, KLK3), prostate cancer markers (PCA3, STEAP1), metabolic regulators (FASN, SOD1), and cell cycle genes (CDKN1A) (Fig. 6D).

Analysis of enriched pathways in the two regions identified T cell and cytokine pathways as upregulated in Region 1 / subclone C2 (Fig. 6E). Co-occurrence analysis showed that cells of the transcriptomic cluster PCa 3 were the most closely associated with CD8+ and exhausted T cells (Fig. 6F). In fact, these T cell subtypes were distributed along the edges of the malignant epithelium in Region 1 (Fig. 6G).

Integration of spatial genomics and spatial transcriptomics in another patient sample (Fig. 6H) also showed mutually exclusive transcriptomic programmes in different subclones as expected (Fig. 6I). However, we also noted that transcriptomic cluster PCa 1 cells span spatially across two malignant subclones (Fig. 6J).

**Figure 6:**
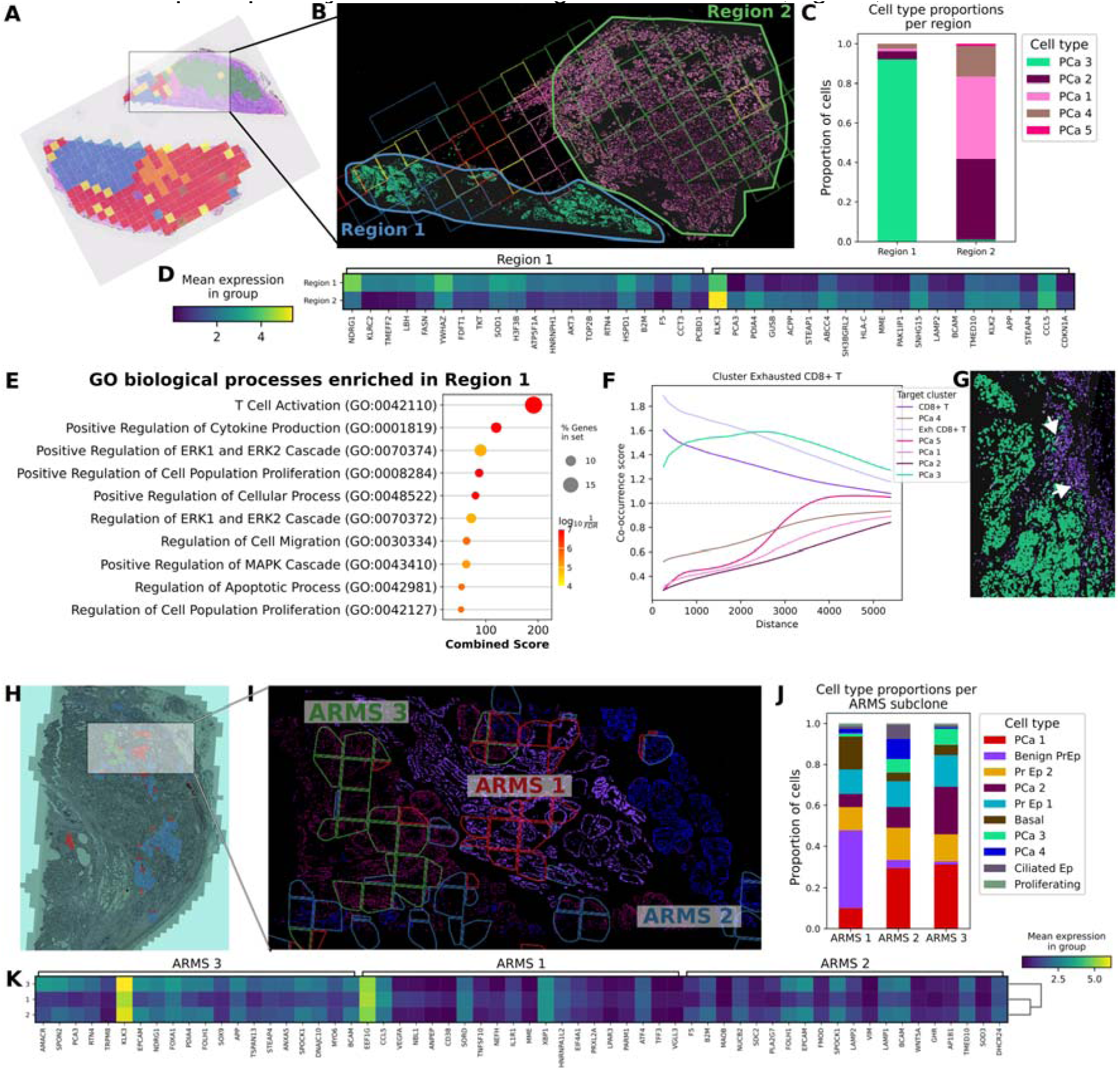
Multi-omic data integration showing subclone-level transcriptomic differences. A) A small portion of tissue (outlined in semi-transparent rectangle) taken from the same block as that used for P15 ARMS DNAseq was assayed using the Xenium Prime spatial transcriptomics platform. B) Region 1 (corresponding to clone C2) and Region 2 (corresponding to clone C3) displayed distinct transcriptomic profiles (cells are coloured by transcriptomic cluster ID). Outlines of the square tiles are coloured by subclone ID. C) Transcriptomic cluster membership between regions 1 and 2 was nearly mutually exclusive. D) Top 20 genes differentially expressed between regions 1 and 2. E) Signalling pathways enriched in Region 1. F) Spatial co-occurrence plot showing enrichment of CD8+ T cells and exhausted T cells close to the PCa 3 transcriptomic cluster (equivalent to clone C2). G) Close-up of Region 1 showing CD8+T cells (purple) and Exhausted T cells (lilac, white arrows) accumulated at the edge of the PCa 3/clone C2 region. H) A small portion of tissue (outlined in semi-transparent rectangle) taken from the same block as that used for 10Q ARMS DNAseq was assayed using the Xenium Prime spatial transcriptomics platform. I) ARMS DNAseq tiles were coloured by subclone IDs and overlaid on spatial transcriptomic data where cells are coloured by transcriptomic cluster ID.

As the Xenium transcriptomic panel included 5100 genes, we sought to obtain inferred copy number alteration calls from these data and test the corroboration with ARMS DNAseq. We used insituCNV, a tool adapted to detect CNAs from imaging-based spatial transcriptomic data^20^. This showed that not all genomic subclones detected by ARMS DNAseq can be identified in the transcriptomics-inferred CNAs (Extended figure E4). For example, in sample P15, losses in chromosomes 10q, 17p, and gains in chromosomes 3, 10p (among others) are detectable through ARMS DNAseq (and confirmed in bulk WGS) but not detected in transcriptomics-inferred CNAs. Conversely, some spurious CNAs (e.g. chromosome 19 gain) were detected in the latter, but were not detected in either ARMS DNAseq or bulk WGS results. However, a few CNAs were seen commonly across both modalities. In P10, focal CNAs seen through ARMS DNAseq are missing in the transcriptomics-inferred CNAs.

**Extended fig E4:**
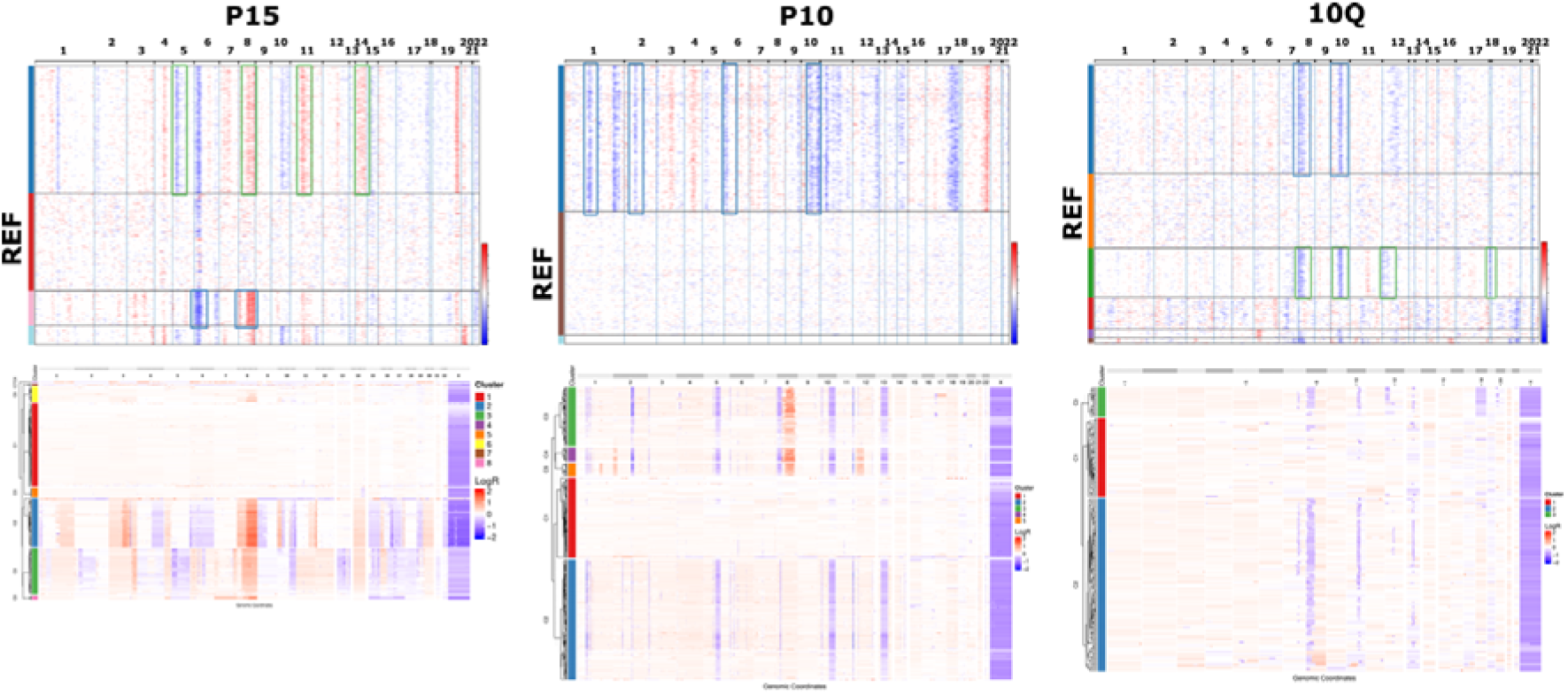
Comparison of CNA calls from transcriptomic inference and ARMS DNAseq. CNAs detected in both transcriptomic inference (top) and ARMS DNAseq (bottom) are highlighted with coloured outline boxes (the colour representing the corresponding cluster ID in the ARMS DNAseq heatmap). REF denotes the transcriptomic cluster that was assigned as the ‘normal’ reference for CNV inference.

**Extended Table ET2:** Comparison between ARMS, bulk WGS, and transcriptomics-inferred CNAs.

| *Patient* | *ARMS tiles* | *ARMS subclones* | *Bulk WGS region* | *Bulk WGS subclones* | *Transcriptomics-inferred subclones* | *Added value of ARMS* |
| --- | --- | --- | --- | --- | --- | --- |
| *P15* | 288 | 4 | 2 | 2 | 3 | 2 additional subclones; Identification of WGD |
| *P10* | 288 | 4 | 8 | 5 | 1 | Better resolution of smaller subclones; missing subclones in ARMS are due to sampling of only one prostate region. |
| *10Q* | 190 | 2 | NA | NA | 2 | Identification of novel subclones; better genomic resolution |

## DISCUSSION

In this study, we describe the ARMS DNAseq method for spatial copy number profiling of cancer samples and demonstrate several applications in histopathology and spatial transcriptomic contexts. Importantly, we show that ARMS DNAseq works well on thin sections of archival FFPE samples, thus unlocking a wealth of material from pathology collections for high resolution spatial genomic analysis. ARMS DNAseq also enables better genomic resolution and subclone identification compared to bulk WGS and copy number inference from spatial transcriptomics. It is important to acknowledge that shallow WGS has been performed at smaller scale, for example in FFPE^21^ or frozen samples^22^. However, we describe here a unified and streamlined workflow that incorporates plate-level barcoding to enable scaling up the analysis to hundreds of spatial regions per experiment. To ease adoption, we have developed software tools to support the ARMS DNAseq workflow, which include 1) a GUI app to enable easy conversion of histopathology annotations to LCM-compatible annotations, 2) a Snakemake workflow to facilitate demultiplexing copy number calling, 3) an R package to call copy number alterations and identify subclones, and 4) a GUI app that enables integration of ARMS DNAseq results with H&E images, Xenium spatial transcriptomics data, and other image data, thus placing the different biological layers in a shared spatial context.

In two of the cases from which we have bulk WGS data (from fresh frozen tissue), ARMS DNAseq analysis of formalin fixed paraffin embedded tissue sections revealed a more detailed subclonal structure and phylogenetic tree compared to the much more expensive bulk WGS approach^10^. In patient #15, we previously showed allele-specific subclonal copy number alterations from two intra-prostatic regions. However, due to complex genomic alterations, we were previously not able to obtain a clean copy number profile from the LPZLat sample using the Battenberg algorithm. We now show, using ARMS DNAseq from a nearby area of the tissue, that we can distinguish an additional subclone (that could not be resolved previously). A combination of the C3 and C8 could potentially explain the (poorly resolved) CNA profile seen in LPZLat bulk WGS. Similarly, in patient #10, ARMS analysis of a single section was sufficient to validate previously identified subclones, and furthermore, resolved smaller (more spatially delimited) subclones compared to bulk WGS. Particularly in this case, the tumour purity of many regions sampled for bulk sequencing was low, limiting its utility. The use of focused genomic analysis and high sampling density, even if shallow, provided more information.

Recent experimental and computational advances in spatial transcriptomics enable the identification of copy number alterations, and consequently subclones, in fresh frozen and FFPE tissue sections. However, the genomic resolution of such methods is limited due to the nature of the assay – when gene expression is being used as a proxy to infer copy number alterations, genomic resolution is a trade-off due to the size of the bins as well as non-coverage of gene-free regions of the genome. Furthermore, spatial transcriptomics-based copy number inference may also be limited by alterations in gene expression as the cancer evolves. In our head-to-head comparison with spatial transcriptomics-inferred copy number profiles from the same samples, ARMS DNAseq achieved greater genomic resolution and reliably identified smaller copy number alterations, which are corroborated by true bulk WGS as well as by pseudobulk analysis of the ARMS data. Indeed, the pseudobulk analysis (either pseudobulking by clone assignments, or based on a *post hoc* hypothesis-driven decision, e.g. tiles belonging to similar morphologies) affords greater confidence to ARMS DNAseq than transcriptomics-inferred results, as a greater genomic resolution is gained due to increased depth per bin. ARMS DNAseq also provides the ground truth and orthogonal confirmation of inferred CNAs.

It is reasonable to assume that at least some morphological diversity is caused by changes in DNA, although changes at the epigenetic, RNA, and protein levels may also be expected to produce morphological heterogeneity without any changes in the DNA upstream of them^23^. In the three cases studied here, tissue regions corresponding to different copy number subclones are often (but not always) morphologically distinct. Conversely, morphologically distinct tissue regions are often genomically distinct in our dataset.

The ability to link morphological subtypes with evolutionary stages of the tumour has implications in a diagnostic and prognostic setting. Evolutionarily ‘late’ morphologies will harbour more mutations and may necessitate a different clinical protocol compared to ‘early’ morphologies. Furthermore, the presence of a diverse range of morphologies within the same patient (morphological heterogeneity) may suggest an associated genomic heterogeneity. This is important in the context of a study by Fernandez-Mateos et al. which showed that genomic and morphological heterogeneity are independent predictors of recurrence in prostate cancer^24^. Sowalsky et al. found that Gleason Pattern 3 and 4 prostate cancer foci were on distinct branches of the phylogenetic tree^25^. Karasaki et al. showed that high grade histological patterns in lung cancer were associated with complex chromosomal changes and later clones^26^. Congruent with these studies, we show through within-patient analysis that later clones harbour more advanced histological subtypes (Gleason patterns 4 and 5, intraductal cribriform) whereas earlier clones from the same lineage have Gleason pattern 3 (an earlier subtype). We also show as a proof of principle, that the genotype-phenotype association can be exploited to predict the latter from the former in samples from the same patient. However, it must be noted that this prediction workflow required scanner-specific batch correction of morphology embeddings and furthermore, limited to within-patient analysis.

Integration of spatial genomics with other spatial modalities can provide information about subclone-specific biology. Co-registration of tissue sections enables the use of a shared coordinate system for such an integration. We used this approach to discover subclone specific distribution of immune cells in one patient, and transcriptional programmes spanning subclone boundaries in another. Further work needs to be done to understand the latter phenomenon – currently it is not known whether this signifies technical artifacts, microenvironmental influences, or plasticity in cell states.

Despite these examples of its utility, the ARMS DNAseq method currently has many limitations. So far, only copy number alterations have been characterised from the ARMS method. Increasing the per-tile depth of sequencing followed by pseudobulk analysis (as performed by Funnell et al. in single cells^27^) may enable the identification of point mutations. Furthermore, as the ARMS library preparation is ‘mini-bulk’ in nature, tumour cell fraction (or purity) is a problem similar to bulk sequencing. This is offset to some extent in solid tumours like prostate cancer; tiles collected from the centre of a tumour mass tend to be relatively pure. However, tiles collected from the edges of a tumour may contain an admixture of malignant and benign cells, resulting in separate clustering of high purity and low purity tiles from the same subclone. Similarly, tiles overlapping the boundaries of two different subclones will have mixed profiles that may appear as noise. Future improvements to address this problem may include more robust exclusion of non-malignant tissue during tile generation, using cell classification foundation models (e.g. Histoplus^28^) to quantify tumour cell percentage in each tile and apply a corresponding purity correction in the copy number calling step, or other improved computational methods. In addition, a laser capture microdissection microscope (which is not widely available) is required for dissecting tissue. Furthermore, unlike spatial transcriptomics methods such as Visium that provide transcriptomic and genomic data, ARMS DNAseq is only suitable for generating genomic data. In cases where sequencing-based spatial transcriptomics is being applied anyway to profile gene expression, obtaining large-scale copy number alteration results at low genomic resolution may be desirable. On the other hand, if a high genomic resolution is required to construct accurate phylogenies over larger areas, ARMS DNAseq is a more suitable approach. ARMS DNAseq, being a direct readout of genomic DNA rather than inference from gene expression data, may also be used for validation of CNAs inferred from spatial transcriptomics.

**Extended Table ET3:**
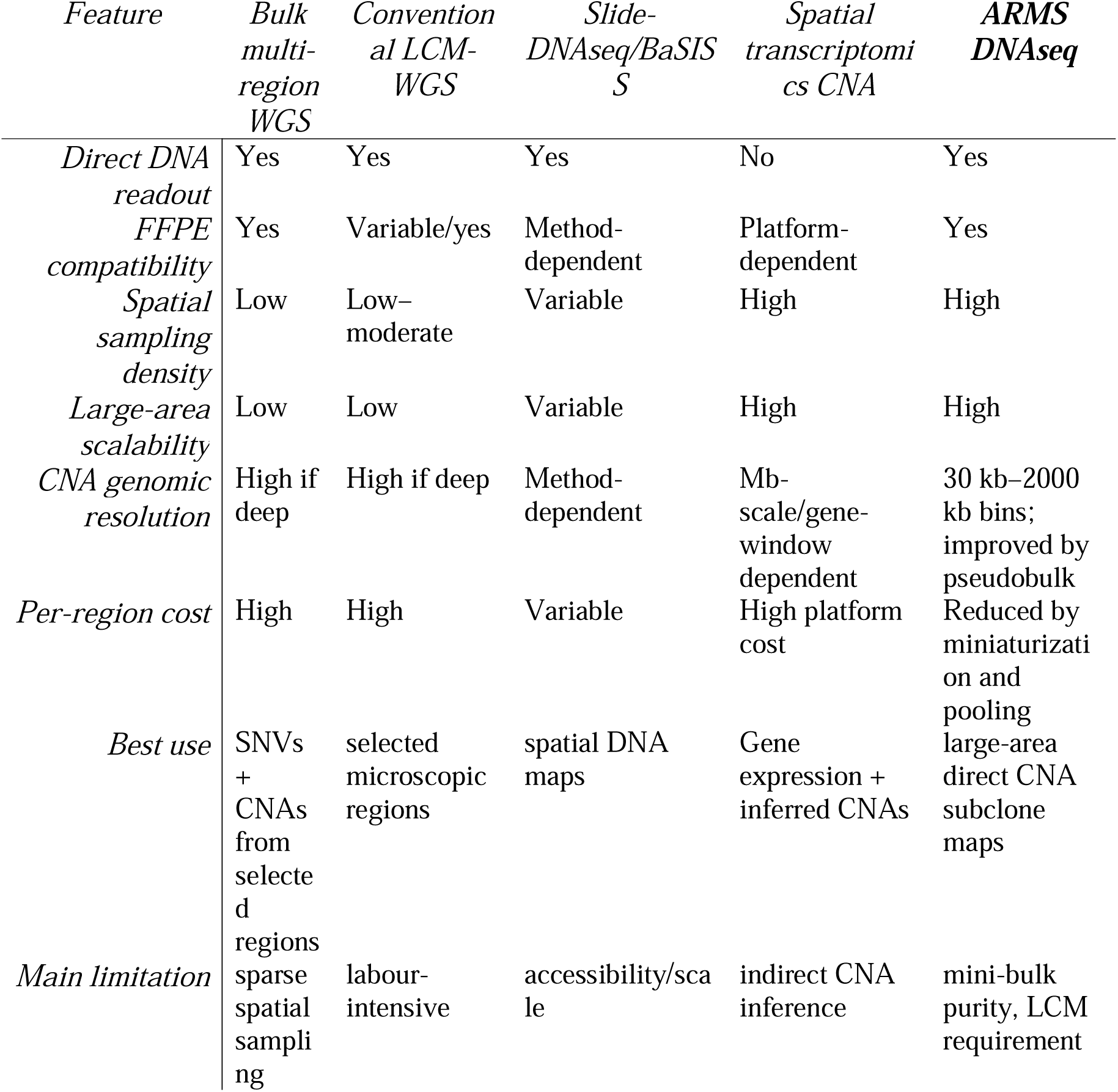
Comparison of advantages and limitations of existing spatial methods with ARMS DNAseq.

In summary, we describe here a scalable method to spatially profile DNA from archival tissue samples, demonstrating its effectiveness in determining copy number alterations from hundreds of regions across three patients. We show use cases for ARMS DNAseq in integrating subclonal architecture with histopathology and spatial transcriptomics, and show its utility in delineating intra-tumour heterogeneity.

## METHODS

### Patient samples

Formalin fixed paraffin embedded tissue samples from the ProMOTE cohort^29^ were obtained from diagnostic archive pathology blocks through the Oxford Radcliffe Biobank, adhering to institutional and national regulations and approved by the ethics committee (REC numbers 24/SC/0220 and 19/SC/0173).

### Slide preparation

Serial formalin fixed paraffin embedded (FFPE) sections were cut on to PEN membrane slides (for LCM) and glass slides (for histopathological assessment). Both types of slides were deparaffinised in Histoclear, rehydrated in descending concentrations of ethanol, and stained with Haematoxylin-Eosin (H&E). Glass slides were coverslipped with DPX mountant and scanned on a Motic slide scanner. PEN membrane slides were temporarily coverslipped with a 1:1 mix of glycerol/water and scanned, following which they were rinsed in water to remove the coverslips, finally washed with 100% ethanol and air dried. PEN slides were stored at 4°C for a few days to weeks until laser capture microdissection.

### Tile annotation

Tissue sections stained with H&E on glass slides were used for determination of region(s) of interest (ROIs) based on features such as histopathological distinctness, benign epithelium, amount of immune cell infiltration, and stromal composition. Such ROIs were then transferred by affine transformation onto the coordinate space of the PEN (LCM) slides. The ROIs were split into square regions of specified size (‘tiles’). The tiles were in turn were divided further into minitiles of no more than 0.3mm x 0.3mm to enable efficient laser capture. Tile annotation can be performed using the ARMS-tiler Napari app https://github.com/rao-genomics-lab/ARMS-tiler.

### Laser capture microdissection

Laser capture microdissection was performed on a Zeiss PALM Microbeam device, with (settings are provided in supplementary information file 1). Tile annotations in whole slide scan coordinates were transferred to LCM coordinates using affine transformation from 3 manually selected points. All functionality from tile generation to transformation of the tiles to the LCM coordinate space is included in the ARMS-tiler Napari app. All minitiles corresponding to a tile were captured into a single well of a 96-well plate, in a total of 40ul lysis buffer. Lysis buffer 1 (30 mM Tris HCl pH 8.0, 0.5% Tween 20, 0.5% NP40, 25 µg/ml Proteinase K) was used for samples P15 and P10 and lysis buffer 2 (0.15% Triton X-100, 0.5 mM EDTA, 25 µg/ml Proteinase K, 1% 2-mercaptoethanol) was used for sample 10Q.

### Library preparation

Samples in lysis buffer were incubated on the lysis program on the thermal cycler (55 °C/15 min, 75 °C/15 min). For library preparation, two versions of the protocol were followed; A) for samples P15 and P10, proteinase K-treated lysates were 1X bead cleaned, followed by combined enzymatic fragmentation and end prep (reaction vol: 4.375 µl; 37 °C/12 min, 65 °C/30 min), adapter ligation (reaction vol: 8.562 µl; 20 °C/20 min), pooling at plate level, bead cleanup, and USER + indexing PCR (reaction vol: 50.5 µl; 20 cycles) B) for sample 10Q, 5 µl proteinase K-treated lysates dried in a 96-well plate (75 °C/30min), followed by end prep (reaction vol: 3.75 µl; 20 °C/30 min, 65 °C/30 min), adapter ligation (reaction vol: 6.1875 µl; 20 °C/20 min), pooling at plate level, bead cleanup, and USER + indexing PCR (reaction vol: 50.5 µl; 20 cycles).

### Sequencing

Sequencing on Illumina NextSeq 500 or NextSeq 6000 with at least 75bp + 75bp paired end sequencing.

### Pre-processing sequencing data

Raw sequencing data were demultiplexed based on i5 and i7 index sequences using bcl2fastq (v2.19.0.316). The demultiplexed fastq (R1 and R2) files represent the combined data from each ligation pool (typically 96 wells), with the first 8 bases on both reads representing the same well-specific barcode followed by an invariant T at position 9. Well-level demultiplexing was performed using Cutadapt (v4.4), thus generating a separate fastq R1 and R2 pair for each tile. Reads were trimmed (fastp), aligned to the human genome (hg38) (bwa mem v0.7.17) and duplicate marked (samtools v1.9) to generate bam files. All preprocessing steps and the respective software package requirements are provided as a Snakemake workflow.

### Copy number calling

ASCAT.sc was used for tile-level copy number calling at a variable bin size (30kb for P15 and P10, 2 Mb for 10Q) and the logR ratio files were used to plot a heatmap of CNAs for each sample. CNA data were clustered using hierarchical agglomerative clustering and visually merged in cases where purity differences resulted in over-clustering. Aligned reads from tiles belonging to a subclone were pseudobulked to generate clone-level CNA plots. All these steps are incorporated into the ARMS R package https://github.com/rao-genomics-lab/ARMS which wraps the ASCAT.sc package^30^ for copy number calling.

### Phylogenetic reconstruction

CNA-based phylogenetic reconstruction was performed manually, based on the principle that descendant clones inherit CNAs from their parent clone. Where two descendent clones harboured a subset of CNAs exclusive to the other, they were placed on separate branches of the phylogenetic tree. Only relative CNAs (logR values) were used; allele-specific information, omitting integer copy number, and WGD information were not used in the reconstruction (although the latter two were derived implicitly through ASCAT.sc analysis of pseudobulked data).

### Feature extraction from whole slide images

Haematoxylin and Eosin (H&E) stained slides were scanned on the Motic or Philips scanner at 40X resolution (pixel size: 0.2627 and 0.22 respectively) or the Hamamatsu scanner for XL slides at 20X resolution (pixel size: 0.44) to yield WSIs in SVS or pyramidal tiff format. Patches of 256x256 pixel size were extracted from the scanned WSIs and provided as input to the Phikon v2 pathology foundation model. Principal component analysis was performed on the resulting 1024-dimension embeddings to reduce the dimensions to 128, followed by hierarchical agglomerative clustering to generate ‘morphology clusters’. The morphology cluster IDs were mapped as distinct colours back to the original tissue coordinates. The Snakemake workflow for the Phikon v2 morphology analysis is available on GitHub https://github.com/rao-genomics-lab/phikon_snakemake_workflow_v2.

### Subclone-morphology association analysis

Each morphology patch was assigned to an ARMS tile by a point-in-polygon test, and it inherited the subclone label of that tile. Patches falling outside every annotated ARMS tile were excluded from all downstream analysis (46,078 of 58,389 patches retained; 78.9%). An association score *z* is calculated as:

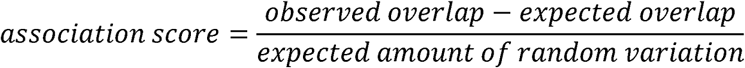

or:

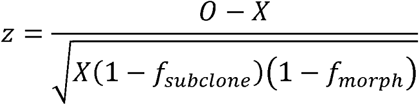

where:

- *0* = observed number of patches with both the chosen subclone and morphology
- *X* = expected number of overlapping patches if the labels were unrelated
- *f_subclone_* = fraction of all patches belonging to that subclone
- *f_morph_* = fraction of all patches belonging to that morphology

The expected overlap is:

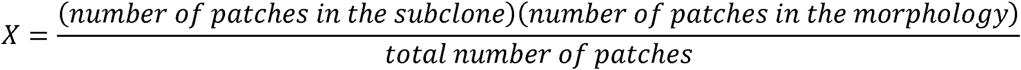

As both nearby morphology clusters and ARMS tiles tended to have the same cluster or subclone IDs (i.e., large areas of tissue had the same morphology or subclone label), a spatially aware null model was deemed necessary to statistically test morphology-subclone association. The genomic (ARMS) field is fixed, while a toroidal shift was applied to the morphology field. 999 permutations of [x, y] offsets were applied, wrapping around at the boundaries. The subclone-morphology association score was recalculated for every shift to produce a null distribution of association scores per subclone-morphology pair. A two-tailed empirical p-value was computed as the proportion of random shifts that produced an association at least as strong as the observed association for that subclone-morphology pair.

The two-tailed empirical p-value is:

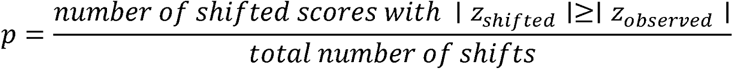

Benjamini-Hochberg False Discovery Rate (FDR) q was applied and both q < 0.05 (denoted by *) and p < 0.05 (denoted by #) are reported in the heatmap.

### Subclone prediction from morphology

Morphology embeddings and ARMS subclone labels from one tissue section were used to train and test a random forest model. Starting with a total of 58,389 patches (20 morphology clusters), clusters (4, 6, 7) were excluded due to low abundance, retaining 46,078 patches (78.9%). Raw (1024-dimension) Phikon embeddings and ARMS subclone labels were used as input, with a 40/10/50 train/validation/test split. The RandomForestClassifier implementation from scikit-learn was used with 200 trees and maximum tree depth of 20. This model was then applied to the same tissue section as well as the remaining sections from the same patient, including a section taken specifically for the Xenium spatial transcriptomics assay. As the sections were scanned on three different scanners (Motic for training slide, Hamamatsu and Philips for inference slides) and the morphology embeddings produced a strong scanner-specific batch effect, a simple centroid translation was applied to the embeddings based on the scanner type, as follows:

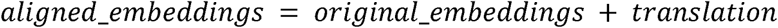

Where,

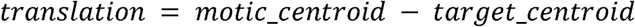

Recall and F1 were low for subclones 5 and 8 potentially due to rarity of these two classes in the training data. Details of the subclone-specific prediction metrics, and section-specific alignment parameters are provided in the supplementary information file.

### Spatial transcriptomic analysis

Serial (within 10s of microns) or near-serial (within 100s of microns) tissue sections from the same blocks as those used for spatial genomics were analysed on the 10X Xenium platform for spatial transcriptomic analysis with a 5000 + 100 (prostate-specific) gene panel. Native Xenium cell segmentation was used, and Leiden clustering (Supplementary information 4, 5) was performed using Scanpy (v1.12). Cell types were annotated manually. Region of interest differential gene expression, spatial neighbourhood and co-occurrence analyses were performed in Scanpy and Squidpy (v1.8).

### Spatial multiomic integration

Grossly recognisable tissue features (e.g. blood vessels, prostate glands, lumina) were used to manually align the spatial genomic and spatial transcriptomic data (and their corresponding H&E images). The spatial transcriptomic assay region was a subset of the spatial genomic assay region. The combined dataset with multiple layers (1. Xenium morphology focus, 2. Xenium cell masks, 3. Xenium nuclear masks, 4. Post-Xenium H&E, 5. ARMS tiles, 6. ARMS H&E, 7. Morphology map) can all be aligned and visualised using the Xenium Viewer https://github.com/sraorao/xenium_viewer which leverages the SpatialData framework^31^ and scverse^32^.

### Copy number inference from spatial transcriptomics

Copy number alterations were inferred from Xenium Prime (5.1K genes) spatial transcriptomics data using the insituCNV package as described by Jensen et al.^20^. Benign luminal epithelial cells were used as the reference for P15 and 10Q. Seminal vesicle epithelial cells were used as the reference for P10, as no benign tissue was present in the Xenium region of interest.

### Code

Claude Sonnet 5 and Opus 5 were used to draft, edit, review, and sanitise Python and R code. All design decisions were manually approved. Some components of the analysis (ARMS DNAseq snakemake workflow, Phikon workflow, parts of the ARMS R package) initially originated as manually generated codebases, which were then further reviewed, and edited using Claude. Detailed changelogs are available in all code repositories generated for this project.

## ACKNOWLEDGEMENTS

This work is funded by a Prostate Cancer UK Research Innovation Award (RIA22-ST2-004) led by SR and CV, and the Cancer Research UK-funded ProMOTE programme grant (C1380/A18444). The Xenium spatial transcriptomic assays were funded by the CRUK Oxford Centre (CTRQQR-2021\100002) as part of the COMBATcancer Prostate Xenium sub-study and the COMBATcancer consortium. We acknowledge the contribution to this study made by the Oxford Centre for Histopathology Research and the Oxford Radcliffe Biobank, which are funded by the University of Oxford, the Oxford CRUK Cancer centre, and the NIHR CRN Thames Valley network.

## DATA AVAILABILITY

Sequencing data is being deposited in the European Genome-Phenome archive and all code used to generate these figures will be deposited in a GitHub repository.

## AUTHOR CONTRIBUTIONS

**C.S.** performed machine learning training and genomic prediction from tissue morphology. **U.S.** analysed spatial transcriptomics data. **A.M.**, **A.W.**, and **M.B.** carried out wet lab experiments (spatial genomics and spatial transcriptomics). **W.B.** and **M.F.** provided feedback on morphology analysis. **S.D.** and **J.Re.** provided laser capture assistance, while **R.F.** and **O.A.** provided laser capture resources. **T.H.** performed genomics data analysis. **S.Mo.**, **R.T.**, and **D.M.-P.** processed tissue samples. **St.M.** and **N.A.** provided computational support. **E.M.** assisted with H&E staining. **D.We.** provided feedback on genomic analysis. **J.H.**, **F.I.**, **C.E.**, **J.Ri.**, **F.H.**, and **D.Wo.** provided resources and feedback. **C.V.** performed pathology analysis, provided feedback, and secured funding. **I.T.**, **R.B.**, and **I.M.** secured grant funding and provided feedback. **S.R.** conducted experimental work, analysed data, secured funding, and wrote the manuscript.

