## Supplementary information for "Scalable spatial DNA sequencing from archival tissue maps copy number subclones"

1. **Workflow times**

Table ST1: Version A workflow time for 192 regions of 1mm^2^

|  | Total time | Hands-on time |
| --- | --- | --- |
| Staining and scanning | 2.5 | 1.0 |
| Affine transformation | 0.5 | 0.5 |
| LCM (1 mm2 90s/tile) – 192 tiles | 5.0 | 1.0 (intermittent) |
| Library prep: lysis | 0.75 | 0.25 |
| Library prep: bead cleanup 1 | 1.0 | 1.0 |
| Library prep: frag + A-tailing | 1.0 | 0.25 |
| Library prep: adapter annealing + ligation | 1.0 | 0.25 |
| Library prep: bead cleanup 2 | 0.75 | 0.25 |
| Library prep: PCR | 1.0 | 0.25 |
| Library prep: adapter cleanup | 3.0 | 1.0 |
| Sequencing | 48 | 1.0 |
|  | 63.5 | 6.75 |

Table ST2: Version A workflow time for 192 regions of 0.10 mm^2^

|  | Total time | Hands-on time |
| --- | --- | --- |
| Staining and scanning | 2.5 | 1.0 |
| Affine transformation | 0.5 | 0.5 |
| LCM (1 mm2 90s/tile) – 192 tiles | 3.2 | 0.5 (intermittent) |
| Library prep: lysis | 0.75 | 0.25 |
| Library prep: bead cleanup 1 | 1.0 | 1.0 |
| Library prep: frag + A-tailing | 1.0 | 0.25 |
| Library prep: adapter annealing + ligation | 1.0 | 0.25 |
| Library prep: bead cleanup 2 | 0.75 | 0.25 |
| Library prep: PCR | 1.0 | 0.25 |
| Library prep: adapter cleanup | 3.0 | 1.0 |
| Sequencing | 48 | 1.0 |
|  | 61.7 | 6.25 |

Table ST3: Version B workflow time for 192 regions of 0.25 mm^2^

|  | Total time | Hands-on time |
| --- | --- | --- |
| Staining and scanning | 2.5 | 1.0 |
| Affine transformation | 0.5 | 0.5 |
| LCM (1 mm2 90s/tile) – 192 tiles | 3.2 | 0.5 (intermittent) |
| Library prep: lysis | 0.75 | 0.25 |
| Library prep: frag + A-tailing | 1.0 | 0.25 |
| Library prep: adapter annealing + ligation | 1.0 | 0.25 |
| Library prep: bead cleanup | 0.75 | 0.25 |
| Library prep: PCR | 1.0 | 0.25 |
| Library prep: adapter cleanup | 3.0 | 1.0 |
| Sequencing | 48 | 1.0 |
|  | 60.7 | 5.25 |

2. **PALM Microbeam laser settings**

[System]

PALMRoboVersion=4.9

PALMControllerVersion=2_Rev.4.2,9162,146M

[Calibration]

Camera=AxioCamICc1

[Calibration_Fluar 5x/0.25 M27]

ReflectorOffset0=0,0

LaserOffset0=580,32

ReflectorOffset1=19,0

LaserOffset1=0,0

ReflectorOffset2=6,0

LaserOffset2=0,0

ReflectorOffset3=0,0

LaserOffset3=0,0

ReflectorOffset4=0,0

LaserOffset4=0,0

ReflectorOffset5=0,0

LaserOffset5=0,0

Calib=0.070374,0.070499

ObjectiveOffset=110,-139

[Calibration_Fluar 10x/0.50 M27]

ReflectorOffset0=0,0

LaserOffset0=268,118

ReflectorOffset1=0,0

LaserOffset1=0,0

ReflectorOffset2=0,0

LaserOffset2=0,0

ReflectorOffset3=0,0

LaserOffset3=0,0

ReflectorOffset4=0,0

LaserOffset4=0,0

ReflectorOffset5=0,0

LaserOffset5=0,0

Calib=0.140192,0.140379

ObjectiveOffset=-110,-268

[Calibration_LD Plan-Neofluar 20x/0.4 Korr M27]

ReflectorOffset0=0,0

LaserOffset0=154,67

ReflectorOffset1=0,0

LaserOffset1=0,0

ReflectorOffset2=0,0

LaserOffset2=0,0

ReflectorOffset3=0,0

LaserOffset3=0,0

ReflectorOffset4=0,0

LaserOffset4=0,0

ReflectorOffset5=0,0

LaserOffset5=0,0

Calib=0.279827,0.280016

ObjectiveOffset=284,-17

[Calibration_LD Plan-Neofluar 40x/0.6 Korr M27]

ReflectorOffset0=0,0

LaserOffset0=77,38

ReflectorOffset1=0,0

LaserOffset1=0,0

ReflectorOffset2=0,-1

LaserOffset2=0,0

ReflectorOffset3=0,-1

LaserOffset3=0,0

ReflectorOffset4=0,0

LaserOffset4=0,0

ReflectorOffset5=0,0

LaserOffset5=0,0

Calib=0.560396,0.561216

ObjectiveOffset=-77,-34

[Calibration_none]

ReflectorOffset0=0,0

LaserOffset0=0,0

ReflectorOffset1=0,0

LaserOffset1=0,0

ReflectorOffset2=0,0

LaserOffset2=0,0

ReflectorOffset3=0,0

LaserOffset3=0,0

ReflectorOffset4=0,0

LaserOffset4=0,0

ReflectorOffset5=0,0

LaserOffset5=0,0

Calib=0.000000,0.000000

ObjectiveOffset=0,0

[Speed]

Stage=220,140,80,40,30,25,

StageDelta=5,5,5,2,2,1,

StageMode=0,0,0,0,0,0,

ModeOnScreen=0,0,0,0,0,0,

Arrow=1149,1500,800,300,200,150,

ArrowDelta=50,50,25,25,10,10,

Cut=150,100,75,46,30,20,

CutDelta=5,5,5,2,1,1,

LPC=1086,1200,600,450,300,200,

LPCDelta=50,50,50,25,25,25,

Position=20000,20000,20000,20000,20000,20000,

PositionDelta=1000,1000,1000,1000,1000,1000,

Scroll=2000,1500,800,300,200,150,

ScrollDelta=50,50,25,25,10,10,

3DTrap=3000,3000,3000,3000,3000,3000,

3DTrapDelta=500,500,500,500,500,500,

[Laser]

CurrentLaserFkt=3

EnergyCut=60,41,41,36,50,50,

EnergyLPC=100,66,65,59,50,50,

EnergyTrap1=50,50,50,50,50,50,

BalanceTrap2=50,50,50,50,50,50,

FocusCut=82,80,65,51,50,50,

FocusLPC=82,82,67,48,50,50,

FocusTrap1=50,50,50,50,50,50,

FocusTrap2=50,50,50,50,50,50,

[AutoChange]

AutoChange=1,42,0,1,25,2,1,24,2,1,23,-3,1,18,-2,1,15,-2,

AutoLPCDistance=40,20,10,6,5,4,

AutoLPCFromLine=20,10,8,3,3,2,

RoboLPCDistance=25,10,7,6,5,4,

DiagonalAutoLPC=1,1,1,1,1,1,

AutoLPCCenterShot=0,0,0,0,0,0,

AutoCircleDistance=100,70,30,12,8,5,

SimultaneousFocus=0,0,0,0,0,0,0,0,0,0,0,0,0,0,0,0,0,0,

[Graphic]

Color=12

ColorCut=9

ColorArea=2

ColorLPC=12

ColorAutoLPCDot=12

ColorRoboLPCCut=8

ColorRoboLPCDot=12

ColorRuler=0

ColorText=0

ColorNrText=15

ThickLine=2

ThickDot=11

ThickRuler=1

[LaserMarker]

ViewLaserMarker=1

ViewTrap1=0

ViewTrap2=0

LaserMarker=0,18,1,30,30,1.000000

LaserMarker3DTrap1=1,6,0,30,30,1.000000

LaserMarker3DTrap2=2,9,0,30,30,1.000000

[Probe]

ImageBeforeCut=0

ImageAfterCut=0

ImageBeforeLPC=0

ImageAfterLPC=0

3. **Organ-level subclone prediction**

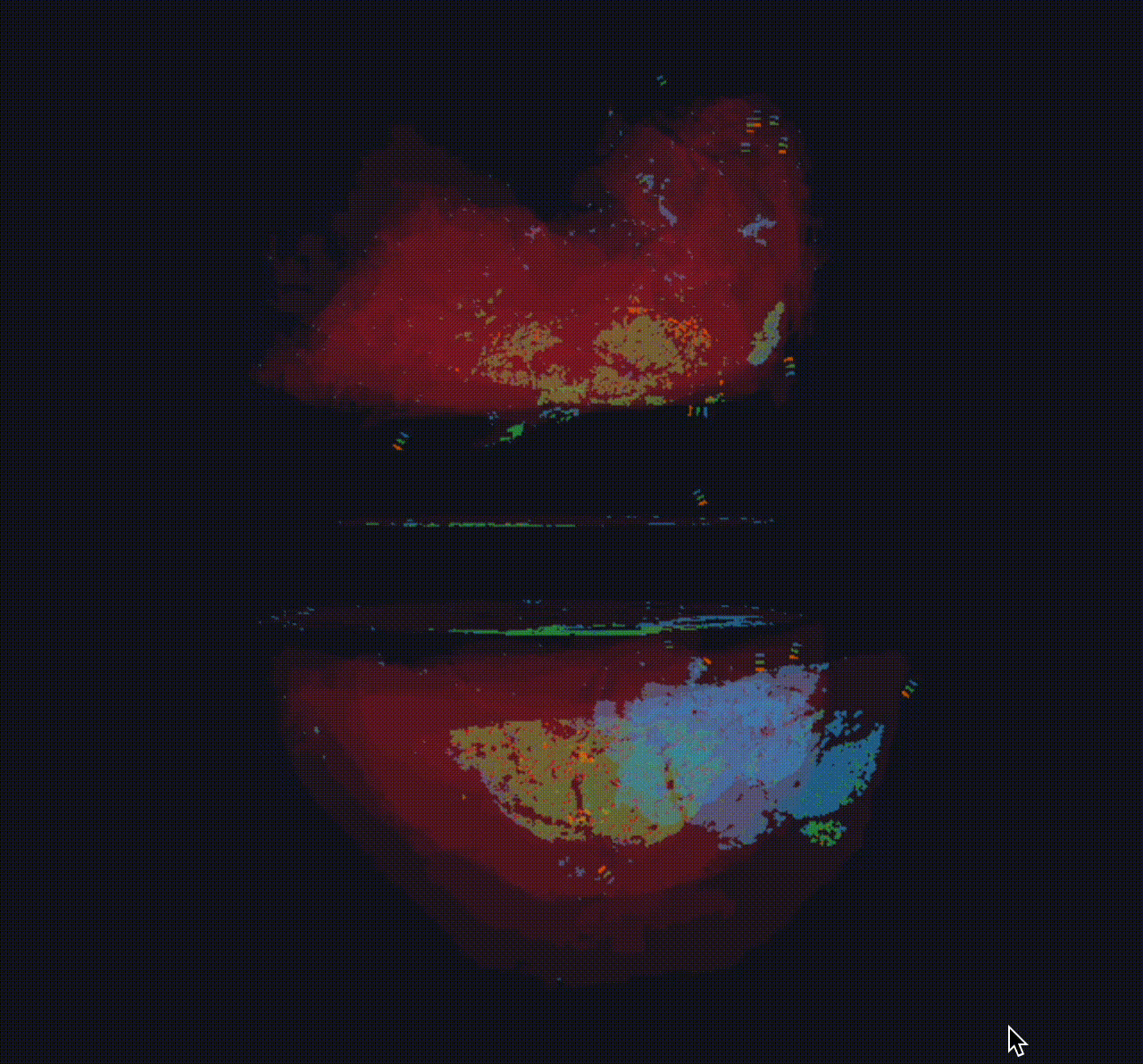

4. **Leiden clustering for P15 Xenium**

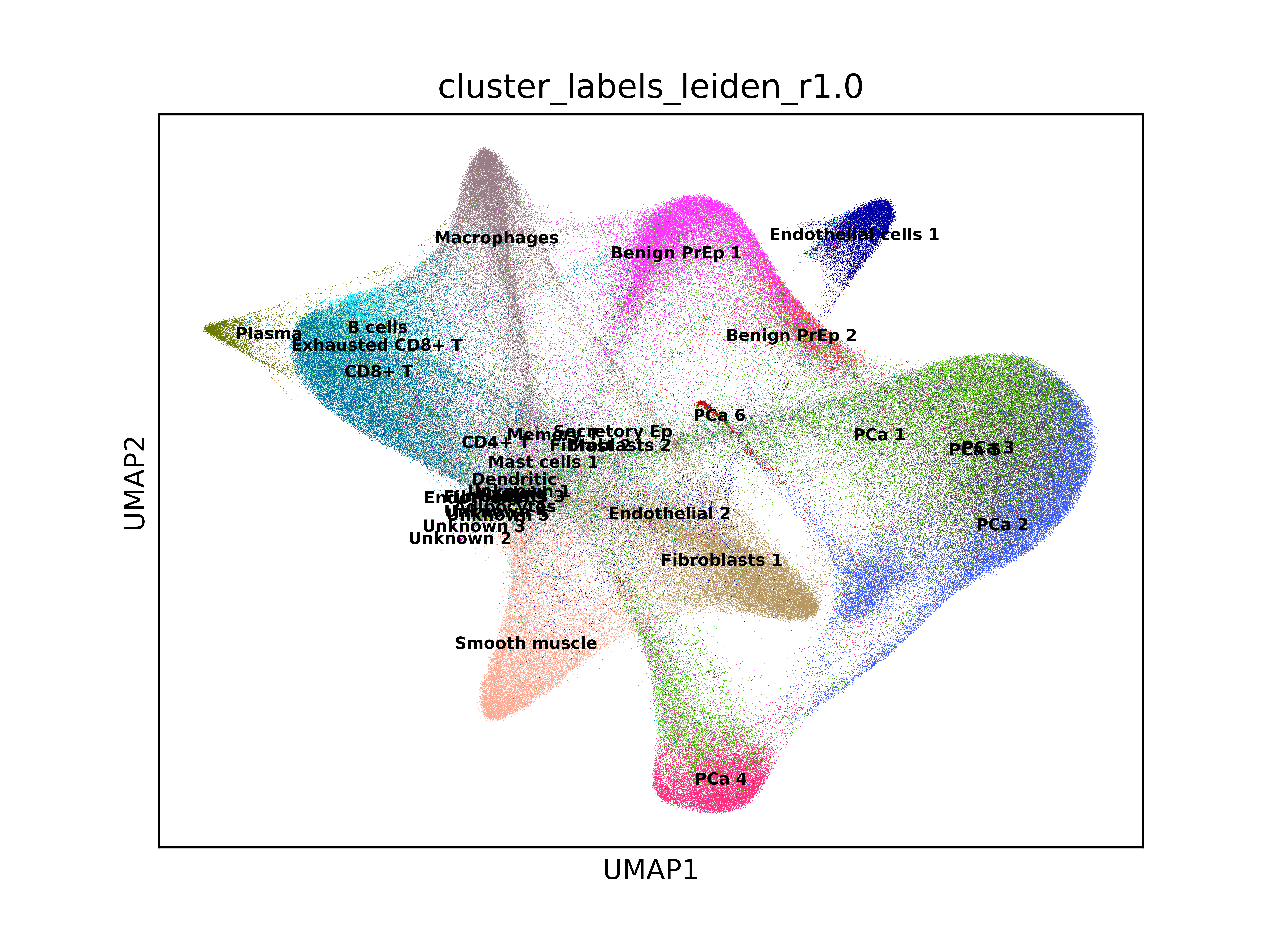

5. **Leiden clustering for 10Q Xenium**

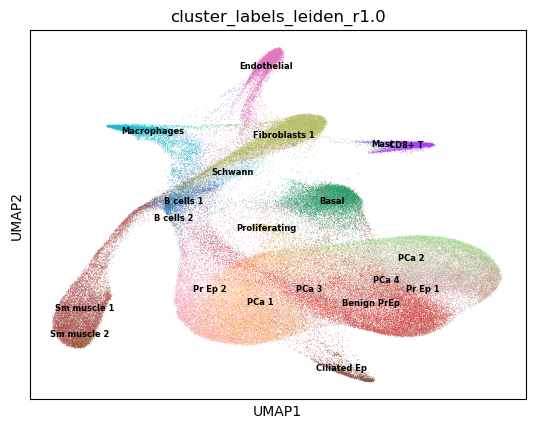
